# Interplay between coding and exonic splicing regulatory sequences

**DOI:** 10.1101/334839

**Authors:** Nicolas Fontrodona, Fabien Aubé, Jean-Baptiste Claude, Hélène Polvèche, Sébastien Lemaire, Léon Charles Tranchevent, Laurent Modolo, Franck Mortreux, Cyril F. Bourgeois, Didier Auboeuf

**Author notes:** Corresponding author: Didier Auboeuf, Laboratory of Biology and Modelling of the Cell, ENS de Lyon, 69007 Lyon, France. Present address: INSERM UMR 861, I-STEM, 28, Rue Henri Desbruères, 91100 Corbeil-Essonnes – France. Equal contribution (Nicolas Fontrodona, Fabien Aubé).

## Abstract

The inclusion of exons during the splicing process depends on the binding of splicing factors to short low-complexity regulatory sequences. The relationship between exonic splicing regulatory sequences and coding sequences is still poorly understood. We demonstrate that exons that are coregulated by any splicing factor share a similar nucleotide composition bias. We next demonstrate that coregulated exons preferentially code for amino acids with similar physicochemical properties because of the non-randomness properties of the genetic code. Indeed, amino acids sharing physicochemical properties correspond to codons that have the same nucleotide composition bias. These observations reveal an unanticipated bidirectional interplay between the physicochemical features encoded by exons and exon splicing regulation by splicing factors. We propose that the splicing regulation of an exon by a splicing factor is tightly interconnected with the physicochemical properties of the exon-encoded protein domain depending on the splicing-factor affinity for specific nucleotides.

## Introduction

Alternative splicing is a cellular process involved in the regulated inclusion or exclusion of exons during the processing of mRNA precursors. Alternative splicing is the rule in human since 95% of human genes produce several splicing variants^1,2^. The exon selection process relies on RNA binding splicing factors that enhance or repress exon inclusion following two main principles. First, splicing factors bind to short intronic or exonic motifs (or splicing regulatory sequences) that are often low-complexity sequences composed of a repetition of the same nucleotide or dinucleotide^3^. The interaction of splicing factors with their cognate binding motifs often depends on the sequence context and on the presence of clusters of related-binding motifs^3–7^. The second principle governing splicing regulation concerns the splicing outcome (i.e., exon inclusion or skipping) that depends on where splicing factors bind on pre-mRNAs with respect to the regulated exons. For example, hnRNP-like splicing factors often repress the inclusion of exons they bind to, but they enhance exon inclusion when they do not bind to exons but instead to their flanking introns^3,8,9^. Meanwhile, exonic binding of SR-like splicing factors (or SRSFs) usually enhances exon inclusion^3,8^.

Since some splicing regulatory sequences lie within protein-coding sequences, a major challenge is to understand how coding sequences accommodate this overlapping layer of information^10–13^. To date, a general assumption is that protein coding regions can accommodate overlapping information or “codes” (including the “splicing code”) as a direct consequence of the redundancy of the genetic code that allows the same amino acid to be encoded by several codons differing only on their third “wobble” nucleotide^10–16^. Therefore, coding and exonic splicing regulatory sequences could evolve independently because of the variation of the third nucleotide of codons. However, it has been shown that some amino acids are preferentially encoded near exon-intron junctions because of the presence of general splicing consensus sequences near splicing sites^17–19^. In addition, recent evidence has suggested that exons that are coregulated in specific pathophysiological conditions may code for protein domains engaged in similar cellular processes ^20,21^. These observations raised the possibility that exons regulated by the same splicing regulatory process code for similar biological information. So far, the lack of large sets of coregulated exons limited studies addressing the interplay between the splicing regulatory process and peptide sequences encoded by splicing regulated exons. By focusing on exons coregulated by different splicing factors, we uncover a bidirectional interplay between the physicochemical protein features encoded by exons and their regulation by splicing factors.

## Results

To investigate the relationship between exonic splicing regulatory sequences and coding sequences, we analyzed RNA-seq datasets generated from different cell lines expressing or not Arg/Ser (RS) domain-containing splicing factors (SRSFs), focusing on SRSF1, SRSF2, SRSF3, and TRA2 (Supplementary Table S1). Each splicing factor regulated a common set of exons in several cell lines but many exons were regulated on a cell line-specific mode (Supplementary Figure S1). It has been shown that SRSF1, SRSF2, SRSF3, and TRA2 bind to GGA-rich motifs^22–25^, SSNG motifs (where S=G or C)^3,23–26^, C-rich and G-poor motifs^24,25,27,28^, and AGAA-like motifs^24,25,29–31^, respectively. As expected, hexanucleotides enriched in SRSF1-, SRSF2-, SRSF3-, or TRA2-activated exons across different cell lines were enriched in purine-rich, S-rich, C-rich, or A-rich motifs, respectively, when compared to control exons and in contrast to exons repressed by the same factor (Figure 1A and Supplementary Table S2).

**Figure 1.**
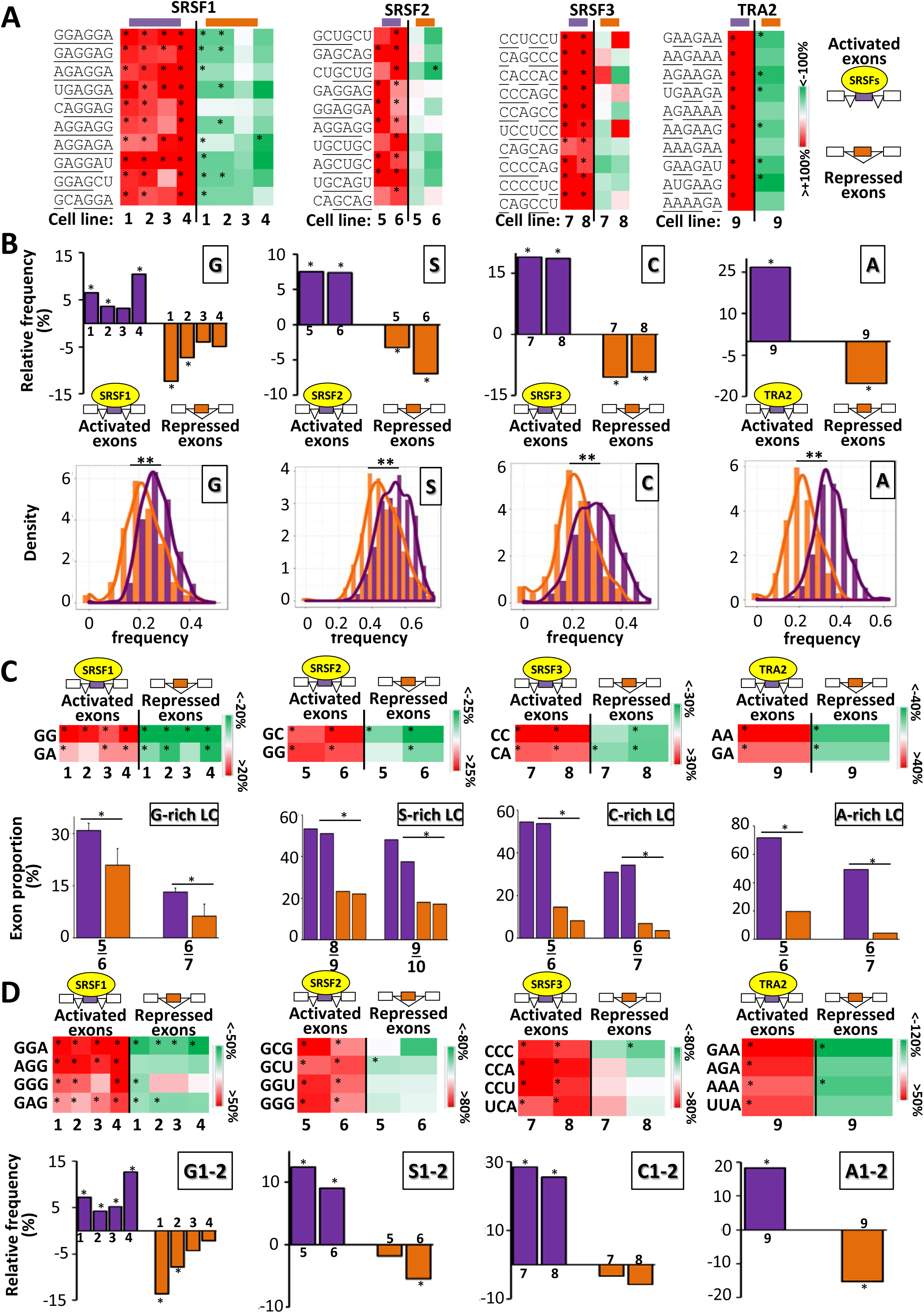
**A**.Color-code of the relative frequency (%) of hexanucleotides in SRSF-activated and -repressed exons. After recovering the 10 most enriched hexanucleotides in SRSF-activated exons, their relative frequency was calculated by comparing it to the average frequency calculated in 10 000 sets of control exons. The relative frequency of these hexanucleotides was also calculated in SRSF-repressed exons. Red and green colors indicate when the frequency is higher and lower, respectively, in the set of regulated exons compared to the sets of control exons. The sets of SRSF-regulated exons originated from publicly available RNAseq datasets generated from the K562 (1), HepG2 (2), GM19238 (3), HeLa (4), K562 (5), Huh7 (6), HepG2 (7), GM19238 (8), and MDA-MB-231 (9) cell lines. Purine, SSNG motifs, Cs, and As are underlined in the enriched hexanucleotides identified in SRSF1-, SRSF2-, SRSF3-, and TRA2-activated exons, respectively. (*) Randomization test FDR-adjusted P-value<0.05 (for the 10 most enriched hexanucleotides). **B**. The upper panels represent the relative frequency (%) when compared to sets of control exons of G, S, C, and A nucleotides in SRSF1-, SRSF2-, SRSF3-, and TRA2-regulated exons, respectively, identified in different cell lines as described in Figure 1A. (*) Randomization test FDR-adjusted P-value <0.05. The lower panels represent the density chart of G, S, C, and A nucleotides in SRSF1-, SRSF2-, SRSF3-, and TRA2-regulated exons, respectively. (**) KS-test <10^−13^. **C**. The upper panels represent the color-code of the relative frequency (%) of dinucleotides, compared to sets of control exons, in SRSF-activated and SRSF-repressed exons across different cell lines as depicted in Figure 1A. The frequency of each dinucleotide was calculated in SRSF-activated and SRSF-repressed exons and expressed as the % of the average frequency calculated in sets of control exons. Red and green colors indicate when the dinucleotide frequency is higher and lower, respectively, in the sets of regulated exons when compared to sets of control exons. Only the two most enriched dinucleotides in SRSF-activated compared to SRSF-repressed exons are represented. (*) Randomization test FDR adjusted P-value<0.05. The lower panels represent the proportion of exons containing at least one low complexity (LC) sequence of 6, 7, 9, or 10 nucleotides. In a sliding window of n nucleotides, the number of the same nucleotide (G, S, C, or A) must be equal to or greater than (n-1). The average of 4 datasets is represented for SRSF1. A logistic regression analysis was performed to test if activated or repressed exons for a given splicing factor have a different content in low complexity sequences while accounting for cell-line variations. (*) P-value<0.05. **D**.The upper panels represent the color-code of the relative frequency (%), compared to sets of control exons, of some codons in SRSF-activated and SRSF-repressed exons across different cell lines as depicted in Figure 1A. The frequency of each codon was calculated in SRSF-activated and SRSF-repressed exons and expressed as the % of the average frequency calculated in sets of control exons. Red and green colors indicate when the codon frequency is higher and lower, respectively, in the sets of regulated exons when compared to sets of control exons. Only some enriched codons identified in SRSF-activated exons are represented. (*) Randomization test FDR-adjusted P-value <0.05. The lower panels represent the relative frequency (%) when compared to sets of control exons, of G (G1-2), S (S1-2), C (C1-2), or A (A1-2) nucleotides at the first and second codon positions in SRSF-activated and -repressed exons identified across different cell lines as depicted in Figure 1A. (*) Randomization test FDR adjusted P-value <0.05.

Each set of factor-specific exons had a specific nucleotide composition bias. Indeed, SRSF1-activated exons were enriched in G when compared to control exons and in contrast to SRSF1-repressed exons (Figure 1B, upper panel and Supplementary Table S2). SRSF2-activated exons were enriched in S (G or C) in contrast to SRSF2-repressed exons (Figure 1B, upper panel). SRSF3-activated exons were enriched in C and impoverished in G (Figure 1B, upper panel and Supplementary Figure S1). Finally, TRA2-activated exons were enriched in A in contrast to TRA2-repressed exons (Figure 1B, upper panel). Accordingly, a larger proportion of SRSF1-, SRSF2-, SRSF3-, or TRA2-activated exons had a high frequency of G, S, C, or A nucleotides, respectively, when compared to the corresponding repressed exons (Figure 1B, lower panels). While the enriched nucleotides within splicing factor-regulated exons could be randomly distributed across exons, we observed an increased frequency of specific dinucleotides and low-complexity sequences. For example, GG, SS, CC, or AA dinucleotides were more frequent in SRSF1-, SRSF2-, SRSF3-, or TRA2-activated exons, respectively, than in control exons or in the corresponding repressed exons (Figure 1C, upper panels and Supplementary Table S2). A larger proportion of SRSF1-, SRSF2-, SRSF3-, or TRA2-activated exons contained G-, S-, C-, or A-rich low-complexity sequences, respectively, when compared to the corresponding repressed exons (Figure 1C, lower panels).

We next analyzed the codon content of exons regulated by these splicing factors. In agreement with the nucleotide composition bias described above, SRSF1-, SRSF2-, SRSF3-, or TRA2-activated exons were enriched in G-, S-, C-, or A-rich codons, respectively, compared to both sets of control exons or the corresponding repressed exons (Figure 1D, upper panels and Supplementary Figure S2 and Table S2). Importantly, the nucleotide composition bias was observed at the first and second codon positions (Figure 1D, lower panels and Supplementary Table S2), raising the possibility that different sets of SRSF-regulated exons may preferentially code for different amino acids.

As shown in Figure 2A (upper panels), amino acids more frequently encoded by SRSF1-, SRSF2-, SRSF3-, and TRA2-activated exons corresponded to G-, S-, C-, and A-rich codons, respectively (see also Supplementary Figure S3 and Supplementary Table S2). This was in sharp contrast to the corresponding repressed exons (Figure 2A, lower panels). For example, glycine (GGN codons) was more frequently encoded by SRSF1-activated exons than by control exons and SRSF1-repressed exons (Figure 2B). A counting of glycine encoded within SRSF1-activated versus SRSF1-repressed exons showed a mirrored distribution: a large proportion of activated exons (∼60%) coded for more than 3 glycine, whereas nearly 70% of repressed exons coded for a maximum of 1 glycine (Figure 2C). Similarly, alanine (GCN codons), proline (CCN codons) and lysine (AAR codons) were more frequently encoded by SRSF2-, SRSF3- and TRA2-activated exons, respectively (Figures 2B and 2C).

**Figure 2.**
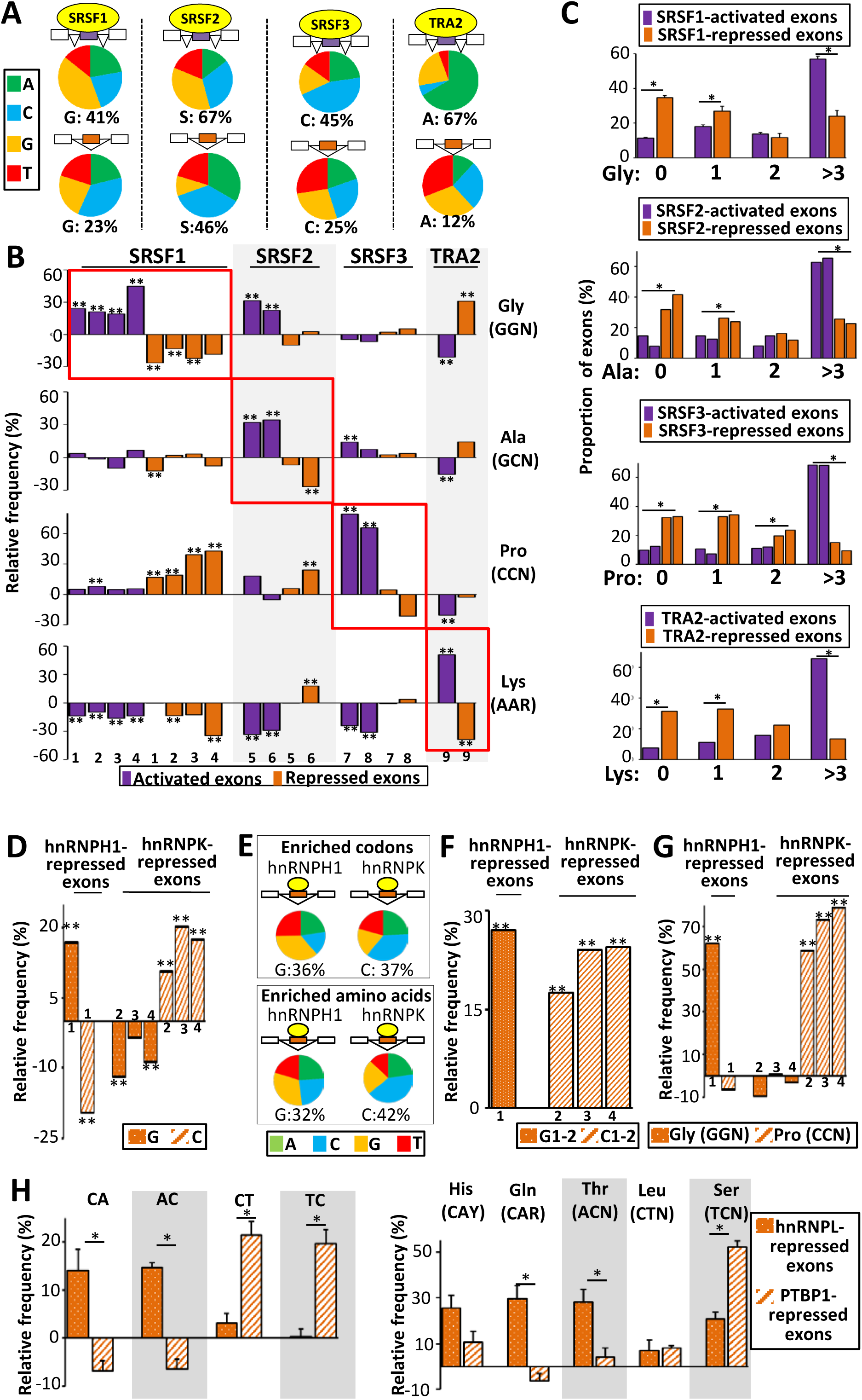
**A**. Nucleotide composition of codons corresponding to amino acids more frequently encoded, when compared to sets of control exons, by SRSF1-, SRSF2-, SRSF3-, or TRA2-activated (upper panels) and - repressed exons (lower panels). **B**.Relative frequency (%) when compared to sets of control exons, of glycine (Gly corresponding to GGN codons), alanine (Ala corresponding to GCN codons), proline (Pro corresponding to CCN codons), and lysine (Lys corresponding to AAR codons) encoded by SRSF1-, SRSF2-, SRSF3-, or TRA2-activated and -repressed exons identified across different cells lines as depicted in Figure 1A. (**) Randomization test FDR adjusted P-value < 0.05. **C**.Proportion (%) of exons from SRSF1-, SRSF2-, SRSF3, or TRA2-regulated exons encoding for 0, 1, 2, and more than three Gly, Ala, Pro, or Lys amino acids, respectively. The average of 4 datasets is represented for SRSF1. A logistic regression analysis was performed to test if activated or repressed exons for a given splicing factor have a different content in codons encoding a particular amino acid while accounting for cell-line variations. (*) P-value<0.05. **D**.Relative frequency (%), when compared to sets of control exons, of G and C nucleotides in hnRNPH1- and hnRNPK-repressed exons identified in 293T (1), K562 (2), GM19238 (3), and HepG2 (4) cell lines. (**) Randomization test FDR-adjusted P-value < 0.05. **E**.Nucleotide composition of the enriched-codons (upper panels) and codons corresponding to enriched amino acids (lower panel) in hnRNPH1- and hnRNPK-repressed exons when compared to sets of control exons. **F**.Relative frequency (%) when compared to sets of control exons, of G (G1-2) or C (C1-2) nucleotides at the first and second codon position from hnRNPH1- or hnRNPK-repressed exons identified in 293T (1), K562 (2), GM19238 (3), and HepG2 (4). (**) Randomization test FDR-adjusted P-value <0.05. **G**. Relative frequency (%) when compared to sets of control exons, of glycine (Gly corresponding to GGN codons) and proline (Pro corresponding to CCN codons) encoded by hnRNPH1- and hnRNPK-repressed exons identified in 293T (1), K562 (2), GM19238 (3), and HepG2 (4) cell lines. (**) Randomization test FDR-adjusted P-value <0.05. **H**.The left panel represents the average of the relative frequency (%) when compared to sets of control exons of CA, CT, AC, and TC dinucleotides calculated from four sets of hnRNPL- or PTBP1-repressed exons. The right panel represents the average of the relative frequency (%) when compared to sets of control exons of histidine (His corresponding to CAY codons), glutamine (Gln corresponding to CAR codons), leucine (Leu corresponding to CTN codons), threonine (Thr corresponding to ACN codons), and serine (Ser corresponding to TCN codons) encoded by four sets of hnRNPL- and hPTBP1repressed exons. (*) Mann-Whitney test P-value<0.05.

We next analyzed several RNA-seq datasets generated from different cell lines expressing or not hnRNP-like RNA-binding proteins that compose a large family of splicing factors that usually repress the inclusion of exons they bind to (Supplementary Table S1 and Supplementary Figure S1). We focused on hnRNPH1, hnRNPK, hnRNPL, and PTBP1 that bind to G-rich motifs^24,25,32^, C-rich motifs^24,25,33^, CA-rich motifs^24,25,27,34^, and CU-rich motifs^24,25,27,35^, respectively. As shown in Figure 2D, hnRNPH1-repressed exons were enriched in Gs, while hnRNPK-repressed exons were enriched in Cs when compared to control exons. Enriched codons in hnRNPH1- and hnRNPK-repressed exons when compared to control exons were enriched in G and C, respectively, and amino acids more frequently encoded by hnRNPH1- and hnRNPK-repressed exons corresponded to G- and C-rich codons, respectively (Figure 2E and Supplementary Figures S2, S3 and Table S2). Very importantly, as in the case of exons activated by SRSF-like splicing factors, the nucleotide composition bias was observed at the first and second codon positions (Figure 2F). Accordingly, glycine (GGN codons) and proline (CCN codons) were more frequently encoded by hnRNPH1-repressed and by hnRNPHK-repressed exons, respectively (Figure 2G).

As mentioned above, hnRNPL represses the inclusion of exons containing CA-rich motifs, while PTBP1 represses the inclusion of exons containing CT-rich motifs^24,25,27,34,35^. CA and AC dinucleotides were enriched in hnRNPL-repressed exons, while CT and TC dinucleotides were enriched in PTBP1-repressed exons compared to control exons (Figures 2H, left panel). Interestingly, histidine (His, CAY codons), glutamines (Gln, CAR codons) and threonine (Thr, ACN codons) were more frequently encoded by hnRNPL-repressed exons than by control exons or PTBP1-repressed exons (Figure 2H, right panel). Meanwhile, serine (Ser, TCN codons) was more frequently encoded by PTBP1-repressed exons than by control exons or hnRNPL-repressed exons (Figure 2H, right panel).

Our results revealed a nucleotide composition bias of splicing factor-regulated exons and a bias regarding the nature of the amino acids that are encoded by these exons. In this setting, it is well established that amino acids sharing similar physicochemical properties (e.g., size, hydropathy, charge) are encoded by similar codons (i.e., composed of the same nucleotides)^36–40^. For example, small amino acids (Ala, Asn, Asp, Cys, Gly, Pro, Ser, Thr), and in particular very small amino acids (Ala, Gly, Ser, Cys) are encoded by S-rich codons while large amino acids (Arg, Ile, Leu, Lys, Met, Phe, Trp, Tyr) are encoded by S-poor codons (Figure 3A). SRSF2 binds to SSNG motifs^3,23–26^ and SRSF2-activated exons are S-rich (see Figure 1). Remarkably, the two sets of analyzed SRSF2-activated exons encoded more frequently for very small and small amino acids when compared to control exons in contrast to SRSF2-repressed exons (Figure 3B and Supplementary Table S2). Conversely, large amino acids were less frequently encoded by SRSF2-activated exons (Figure 3B). Accordingly, a larger proportion of SRSF2-activated exons corresponded to peptides containing very small amino acids, and a smaller proportion of these exons corresponded to peptides containing large amino acids, when compared to SRSF2-repressed exons (Figure 3C).

**Figure 3.**
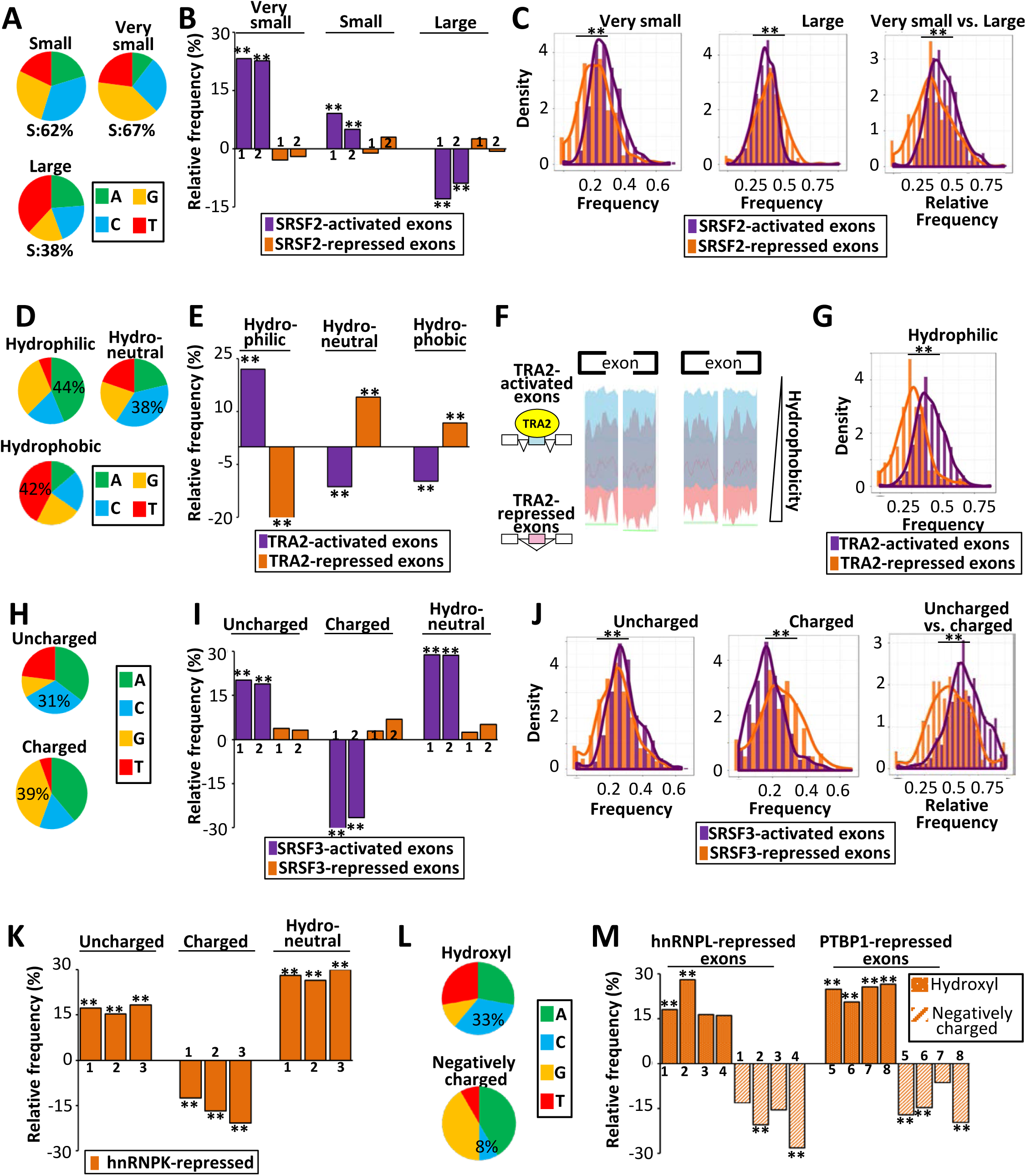
**A**.Nucleotide composition of codons encoding small, very small and large amino acids. S=G or C. **B**.Relative frequency (%) when compared to sets of control exons, of very small, small and large amino acids encoded by two sets of SRSF2-activated and -repressed exons identified in the K562 (1) and Huh7 (2) cell lines. (**) Randomization test FDR-adjusted P-value <0.05. **C**.Density chart of SRSF2-activated and -repressed exons identified in K562 cells coding for very small and large amino acids. (**) KS-test <10^−5^. **D**.Nucleotide composition of codons encoding hydrophobic, neutral and hydrophilic amino acids. **E**.Relative frequency (%) when compared to sets of control exons, of hydrophilic, neutral and hydrophobic amino acids encoded by TRA2-activated and TRA2-repressed exons. (**) Randomization test FDR-adjusted P-value <0.05. **F**.Hydrophobic scales of TRA2-activated and TRA2-repressed exons. The green bottom line indicates the Mann-Whitney test P-value <0.05 at each amino acid position. **G**.Density chart of TRA2-activated or –repressed exons coding for hydrophilic amino acids. (**) KS-test <10^−13^. **H**. Nucleotide composition of codons encoding polar uncharged and charged amino acids. **I**.Relative frequency (%) when compared to sets of control exons, of polar uncharged, charged or neutral (in terms of hydropathy) amino acids encoded by two sets of SRSF3-activated and -repressed exons identified from HepG2 (1) and GM19238 (2) cell lines. (**) Randomization test FDR-adjusted P-value <0.05. **J**.Density chart of SRSF3-activated and -repressed exons coding for uncharged and charged amino acids. (**) KS-test <10^−4^. **K**.Relative frequency (%) when compared to sets of control exons, of polar uncharged, charged or neutral (in terms of hydropathy) amino acids encoded by three sets of hnRNPK-repressed exons identified from K562 (1), GM19238 (2), and HepG2 (3) cell lines. (**) Randomization test FDR-adjusted P-value <0.05. **L**.Nucleotide composition of codons encoding for hydroxyl-containing and negatively charged amino acids. **M**.Relative frequency (%) when compared to sets of control exons, of hydroxyl-containing and negatively charged amino acids encoded by four sets of hnRNPL-repressed exons (K562 (1), HepG2 (2), LNCaP (3), GM19238 (4)) and four sets of PTBP1-repressed exons (HepG2 (5), 293T (6), HeLA (7), K562 (8) cells). (**) Randomization test FDR-adjusted P-value <0.05.

Amino acids can be classified in three families in regards to their hydropathy and each family is encoded by codons having different features. Hydrophilic amino acid (Arg, Asn, Asp, Gln, Glu, Lys) are encoded by A-rich codons, hydrophobic amino acids (Ala, Cys, Ile, Leu, Met, Phe, Val) are encoded by T-rich and A-poor codons, and ambivalent or neutral amino acids (Gly, His, Pro, Ser, Thr, Tyr) are encoded by C-rich codons^37,39,41–45^ (Figure 3D). TRA2 that binds to AGAA-like motifs activates the inclusion of A-rich exons (see Figure 1). Interestingly, TRA2-activated exons encoded hydrophilic amino acids more frequently and neutral or hydrophobic amino acids less frequently than TRA2-repressed or control exons (Figure 3E and Supplementary Table S2). This result was confirmed using different hydrophobicity propensity scales (Figure 3F). Accordingly, a larger proportion of TRA2-activated exons compared to TRA2-repressed exons encoded peptides containing hydrophilic amino acids (Figure 3G).

Polar uncharged amino acids (Asn, Gln, Ser, Thr, Tyr) correspond to C-rich and G-poor codons, while polar charged amino acids (Asp, Glu, Lys, Arg) correspond to G-rich and C-poor codons^37,42^ (Figure 3H). As described above, SRSF3 binds to C-rich motifs and activates exons that are C-rich and G-poor (see Figure 1). Remarkably, the two sets of SRSF3-activated exons encoded uncharged amino acids more frequently than control exons or SRSF3-repressed exons and vice-versa for polar uncharged amino acids (Figure 3I and Supplementary Table S2). They also encoded more frequently hydropathically neutral amino acids (Figure 3I) that correspond to C-rich codons (Figure 3D). As shown in Figure 3J, a larger proportion of SRSF3-activated exons (compared to SRSF3-repressed exons) corresponded to peptides composed of polar uncharged amino acids, while a smaller proportion of these exons corresponded to peptides composed of polar charged amino acids. Similar observations were made for three independent sets of exons repressed by hnRNPK (Figure 3K) that represses C-rich exons (see Figure 2).

Hydroxyl-containing amino acids, including serine and threonine, become negatively charged when phosphorylated. These amino acids are encoded by C-rich codons, while negatively charged amino acids are encoded by C-poor codons (Figure 3L). Remarkably, exons repressed by PTBP1 and hnRNPL (respectively CT- and CA-rich exons, see Figure 2) encoded hydroxyl-containing amino acids more frequently than control exons (Figure 3M and Supplementary Table S2). Meanwhile, they encoded negatively charged amino acids less frequently (Figure 3M).

The physicochemical properties of amino acids are often more or less related. For example, hydrophilic amino acids are often charged amino acids. We observed that exons regulated by a given splicing factor often coded for amino acids that have different physicochemical properties depending on whether the exons are activated or repressed by this factor (Figure 3). Consequently, each splicing factor induced a shift toward different combinations of protein physicochemical properties encoded by their regulated exons (Figure 4A). For example, SRSF2 favored the inclusion of exons encoding very small amino acids rather than large amino acids, in contrast to TRA2 (Figures 4B and 4C). Conversely, TRA2 favored the inclusion of exons encoding hydrophilic amino acids rather than hydrophobic ones, in contrast to SRSF2 (Figures 4B and 4C). Likewise, SRSF3 favored the usage of exons encoding uncharged protein regions rather than charged regions, in contrast to SRSF1 that favored the inclusion of exons encoding charged amino acids (Figures 4B and 4D).

**Figure 4.**
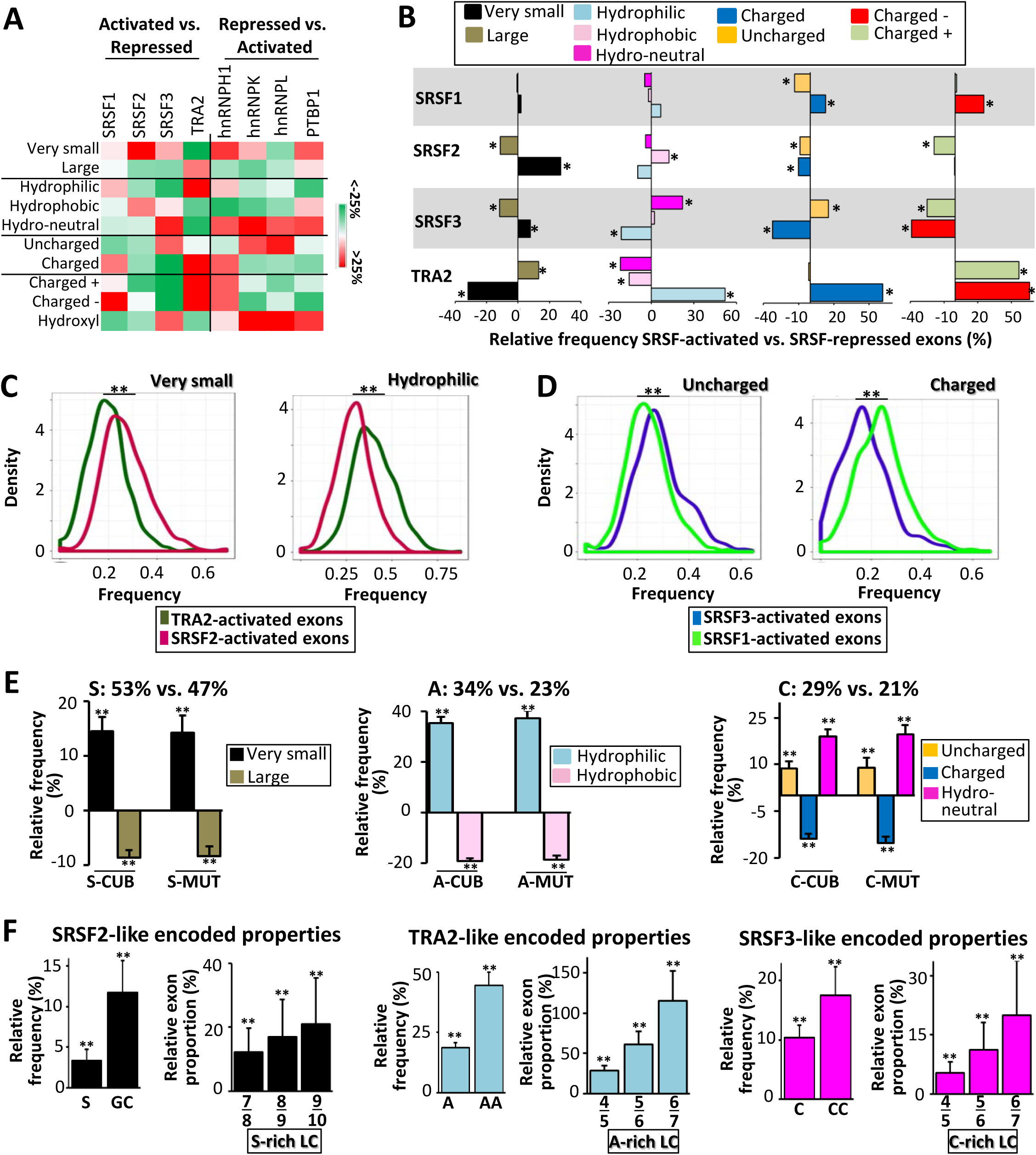
**A**.Color-code corresponding to the relative frequency (%) of amino acid physicochemical properties as indicated when comparing all the SRSF1-, SRSF2-, SRSF3-, and TRA2-activated exons to all the SRSF1-, SRSF2-, SRSF3-, and TRA2-repressed exons, respectively, or when comparing all the hnRNPH1, hnRNPK, hnRNPL, and PTBP1-repressed exons to all the hnRNPH1, hnRNPK, hnRNPL, and PTBP1-activated exons, respectively. **B**.Relative frequency (%) of very small, large, hydrophilic, neutral, hydrophobic, charged, uncharged, negatively charged (charged -), and positively charged (charged +) amino acids when comparing all the SRSF1-, SRSF2-, SRSF3-, or TRA2-activated exons to all the SRSF1-, SRSF2-, SRSF3-, or TRA2-repressed exons, respectively. (*) Mann-Whitney FDR-adjusted p-value <0.05. **C**.Density chart of TRA2-activated and SRSF2-activated exons coding for very small or hydrophilic amino acids. (**) KS-test <10^−13^. **D**.Density chart of SRSF3-activated and SRSF1–activated exons coding for polar uncharged or charged amino acids. (**) KS-test <10^−13^. **E**.The left panel represents the relative frequency (%) of very small and large amino acids encoded by 100 sets of 300 generated-exonic sequences containing a high frequency (53%) of the S nucleotide (S-CUB and S-MUT) compared to 100 sets of 300 generated-exonic sequences containing a low S-nucleotide frequency (47%). The middle panel represents the relative frequency (%) of hydrophilic and hydrophobic amino acids encoded by exonic sequences containing a high frequency (34%) of A nucleotide (A-CUB and A-MUT) compared to exonic sequences containing a low A-nucleotide frequency (23%). The right panel represents the relative frequency (%) of polar uncharged, charged and neutral (in terms of hydropathy) amino acids encoded by exonic sequences containing a high frequency (29%) of Cs(C-CUB and C-MUT) compared to exonic sequences containing a low C-nucleotide frequency (21%). (**) t-test p-value <10^−30^. **F**.The left panels represent the relative frequency (%) of the S nucleotide and GC dinucleotide as well as the relative proportion (%) of exons with S-rich low-complexity (LC) sequences of 100 sets of 300 mutated exons encoding for the same physicochemical properties than SRSF2-activated exons compared to 100 sets of 300 mutated exons encoding for the same physicochemical properties than SRSF2-repressed exons. The middle panels represent the relative frequency (%) of the A nucleotide and AA dinucleotide as well as the relative proportion (%) of exons with A-rich low-complexity sequences of mutated exons encoding for the same physicochemical properties than TRA2-activated exons compared to mutated exons encoding for the same physicochemical properties than TRA2-repressed exons. The right panels represent the relative frequency (%) of the C nucleotide and CC dinucleotide as well as the relative proportion (%) of exons with C-rich low-complexity (LC) sequences of mutated exons encoding for the same physicochemical properties than SRSF3-activated exons compared to mutated exons encoding for the same physicochemical properties than SRSF3-repressed exons. (**) t-test p-value <10^−30^.

On one hand, splicing factors bind to sequences that have a biased nucleotide composition and on the other hand, amino acids with similar physicochemical properties are encoded by codons having the same nucleotide composition bias. Therefore, we anticipated that increasing the exonic density of specific nucleotides as measured in splicing factor-regulated exons would increase the density of encoded amino acids sharing the same physicochemical properties, as observed in those exons. To challenge this possibility, we generated random exonic coding sequences enriched in specific nucleotide(s) by following the human codon usage bias (labelled CUB sequences) or by randomly mutating human coding exons (labelled MUT sequences). For example, we generated 100 sets of 300 coding exons containing either 53% or 47% of S nucleotides, as measured in SRSF2-activated and SRSF2-repressed exons, respectively. Increasing by ∼13% the density of S nucleotides in coding exons (S-CUB or S-MUT) increased (by ∼15%) the frequency of encoded very small amino acids, while it decreased (by ∼10%) the frequency of encoded large amino acids (Figure 4E, “S=53% vs 47%”), as observed for SRSF2-activated and SRSF2-repressed exons (Figure 3B). The increase of the density of A nucleotides in coding exons (A-CUB and A-MUT) from 23% to 34%, as measured in TRA2-repressed and TRA2-activated exons, respectively, increased (by ∼40%) the frequency of encoded hydrophilic amino acids and it decreased (by ∼20%) the frequency of encoded hydrophobic amino acids (Figure 4E, “A= 34% vs 23%”), as observed for TRA2-activated and TRA2-repressed exons (Figure 3E). Finally, an increase in the density of C nucleotides in coding exons (C-CUB and C-MUT) from 21% to 29%, as measured in SRSF3-repressed and SRSF3-activated exons, respectively, increased the frequency of encoded uncharged amino acids and neutral amino acids while it decreased the frequency of encoded charged amino acids (Figure 4E, “C= 29% vs 21%”), as observed for SRSF3-activated and SRSF3-repressed exons (Figure 3I).

We next generated exons coding for different proportions of amino acids sharing the same physicochemical features by mutating randomly-selected human coding exons. For example, we generated 100 sets of 300 mutated exons encoding different proportions of very small and large amino acids, using their respective frequency measured in SRSF2-activated or SRSF2-repressed exons (See Materials and Methods). Mutated exons coding more frequently very small rather than large amino acids had a higher frequency of S nucleotides and GC dinucleotides and contained more frequently S-rich low complexity sequences (Figure 4F, “SRSF2-like encoded properties”), as observed when comparing SRSF2-activated to SRSF2-repressed exons (Figure 1). We next generated exons preferentially encoding hydrophilic rather than hydrophobic amino acids by using their respective frequency measured in TRA2-activated or TRA2-repressed exons, respectively. Mutated exons encoding more frequently hydrophilic amino acids had a higher frequency of A nucleotides and AA dinucleotides and contained more frequently A-rich low complexity sequences (Figure 4F, “TRA2-like encoded properties”), as observed when comparing TRA2-activated to TRA2-repressed exons (Figure 1). Finally, we generated exons preferentially encoding hydropathically neutral rather than charged amino acids, using their respective frequency measured in SRSF3-activated or SRSF3-repressed exons, respectively. Mutated exons encoding more frequently hydropathically neutral amino acids had a higher frequency of C nucleotides and CC dinucleotides and contained more frequently C-rich low complexity sequences (Figure 4F, “SRSF3-like encoded properties”), as observed when comparing SRSF3-activated to SRSF3-repressed exons (Figure 1).

Altogether, these results demonstrate that increasing the exonic density of specific nucleotides based on the frequencies observed in specific sets of splicing factor-coregulated exons, increased the frequency of encoded amino acids having specific physicochemical properties, as observed for the exons coregulated by the corresponding splicing factor. Conversely, increasing the density of exon-encoded amino acids with specific physicochemical properties based on the frequencies observed in specific sets of splicing factor-coregulated exons, increased the exonic frequency of specific nucleotides and low complexity nucleotide sequences that correspond to motifs recognized by the corresponding splicing factor.

## Discussion

Sets of exons coregulated by a given splicing factor are enriched for specific low-complexity sequences (often composed of a repeated (di)nucleotide) that correspond to their cognate RNA binding sites (Figures 1A and 1C). We showed that each set of exons of which inclusion (or exclusion) is enhanced by a given splicing factor is enriched for specific nucleotide(s) when compared to control exons or to exons repressed (or activated, respectively) by the same splicing factor (Figures 1B, 2D, and 2H). To the best of our knowledge, this splicing-related exonic nucleotide composition bias has not been reported yet. However, it is in agreement with recent observations indicating that the interaction of a splicing factor with a binding motif depends on the sequence context and on the presence of clusters of related binding motifs ^3–7^. For example, increasing the exonic frequency of GGA-like motifs increases the probability of an exon to be regulated by the SRSF1 splicing factor that binds to GGAGGA-like motifs even though only one binding site is used ^6^.

Coding sequences overlap several kinds of regulatory sequences, including exonic splicing regulatory sequences. To date, it has been assumed that the redundancy of the genetic code permits protein-coding regions to carry this extra information^10–16^. This means that the sequence constraints imposed by splicing factor binding motifs would accommodate with coding sequences by impacting only the third codon position. However, we observed that nucleotide composition bias of splicing factor-regulated exons impacts the first and second codon positions (Figures 1D and 2F). Since amino acids having the same physicochemical properties correspond to codons with similar nucleotide composition bias, a direct consequence of the exonic nucleotide composition bias associated with the splicing regulatory process is that each set of splicing factor-regulated exons preferentially encodes amino acids having similar physicochemical properties (Figures 2B, 2G, 2H, and 3). Therefore, we propose that the interplay between coding and exonic splicing regulatory sequences that we report is based on a straightforward principle related to both the non-randomness of the genetic code and the preferential binding of splicing factors to low-complexity sequences. Because of these properties, the high exonic density of a specific nucleotide related to splicing factor-binding features increases the probability that an exon encodes amino acids with similar physicochemical properties (Figure 4E). Conversely, the high density of amino acids corresponding to specific physicochemical properties increases the probability of generating exonic nucleotide composition bias and nucleotide low complexity sequences (Figure 4F). In this setting, the function of splicing factors would not only be to regulate the production of individual specialized protein isoforms, but it would also be to control more globally the intracellular content of specific protein domains having specific physicochemical properties. Each splicing factor would control a specific combination of exon-encoded protein physicochemical properties accordingly to its affinity for specific nucleotides.

## Materials and Methods

### RNA-seq dataset analyses

Publicly available RNA-seq datasets generated from different cell lines expressing or not various splicing factors as described in Supplementary Table S1 were analyzed using FARLINE. FARLINE is a computational program dedicated to analyze and quantify alternative splicing variations, as previously reported^46^.

### Frequency of hexanucleotides, dinucleotides, nucleotides, codons, amino acids, and amino acid physicochemical features in exon sets

The formula (1) was used to compute the frequencies of words (*D*_*n*_) of size *n* within a set of exons *S*_*n*_ *=* {*y*_*1*_, *…, y*_*N*_} such that *y*_*i*_ is an exon *i* having a number *L*_*i*_ of nucleotides:

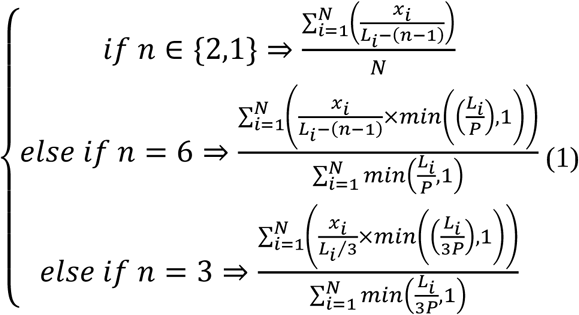

where *x*_*i*_ is the number of occurrences of *D*_*n*_ in exon *i, n* is set to 6, 2, and 1 for hexanucleotides, dinucleotides and nucleotides, respectively. For codons, amino acids and amino acid physicochemical properties, n is set to 3. *P=*50 is a penalty size used to decrease the border effects seen in small exons and *N is* the number of exons in the set *S*_*n*_. For hexanucleotides and dinucleotides, the occurrences x_i_ of *D*_*n*_ are overlapping whereas they are contiguous for the others. In the particular case of amino acids and amino physiochemical properties *D*_*n*_ represents a group of codons encoding the same amino acid or the same physiochemical properties respectively. When coding phase is mandatory, incomplete codons at exon borders were deleted.

Very small (Ala, Gly,Ser, Cys), small (Ala, Asn, Asp, Cys, Gly, Pro, Ser, Thr), large (Arg, Ile, Leu, Lys, Met, Phe, Trp, Tyr), polar uncharged (Asn, Gln, Ser, Thr, Tyr), charged (Asp, Glu, Lys, Arg), hydroxyl-containing (Ser, Thr, Tyr), hydrophilic (Arg, Asn, Asp, Gln, Glu, Lys), hydro-neutral (Gly, His, Pro, Ser, Thr, Tyr), and hydrophobic (Ala, Cys, Ile, Leu, Met, Phe, Val) amino acids were classified as previously reported (*43-45*).

The hydrophobicity scale was calculated as defined by Kyte et al.(*44*) and Engelman et al(*45*). TRA2-activated or -repressed exons larger or equal than 30 amino acids were selected to calculate the average of hydrophobicity using a sliding window of 5 amino acids with a step of 1 amino acid for the 30 first and 30 last amino acids. The mean and the standard deviation of the hydrophobicity values corresponding to each exon set were then calculated for each position of the window.

### Generation of sets of control exons and statistical analyses

To test whether a feature was enriched in a set *S*_*N*_ of *N* exons, a randomization test was made by sampling, from FasterDB^47^, 10000 sets of control exons, **C** = {C_1_, …, C_10000_}, with C_l_ = { *y*_*l,1*_, *…, y*_*l,i*_} such that *y*_*l,i*_ is the exon *i* having a number of *L*_*l,i*_ nucleotides following the constraints:

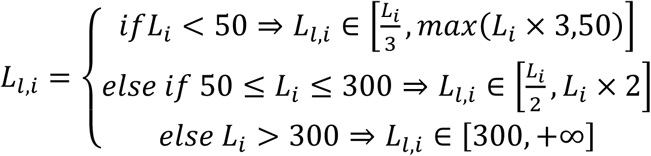

where 50 and 300 nucleotides correspond to the 5^th^ and 95^th^ quantile of the distribution of the length of all the exons defined in FasterDB.

The relative frequency of a feature *D*_*n*_ in *S*_*n*_ compared to the sets of control exons **C** was calculated by the formula:

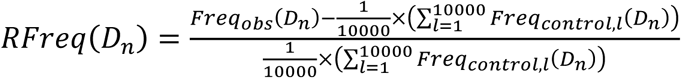

Where *Freq*_*obs*_(*D*_*n*_) is the frequency (as in equation (1)) of a word *D*_*n*_ of size *n* in *S*_*n*_ and 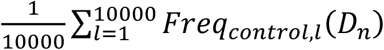 is the average frequency (as in equation (1)) of *D*_*n*_ in **C**.

In order to calculate an empirical p-value, the number of control frequencies upper or lower than the frequency in the set of interest is determined. Then the smallest number between those two is kept and divided by the number of control sets (*i.e.,* 10000).

All p-values obtained for each set of features have been corrected using Benjamini-Hochberg correction.

The nucleotide composition of enriched codons or codons corresponding to enriched amino acids was calculated after recovering codons or amino acids whose frequency was 10% higher in the set of exons of interest than their average frequency in sets of control exons.

### Low complexity sequences and random sequences

Low complexity sequences were defined as sequences of n (n=5 to 10) nucleotides containing at least (n-1) occurrences of the same nucleotide.

Random exonic sequences (from 50- to 300-long nucleotides) with specific nucleotide composition bias were generated using two strategies. First, random codons sequences respecting the human codon usage bias (CUB exons) were generated. These sequences were then mutated randomly, one nucleotide at a time, to increase or decrease the frequency of a specific nucleotide.

Only mutation increasing or decreasing the frequency toward 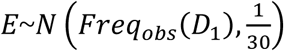 (where *Freq* _*obs*_(*D*_1_) is the nucleotide frequency observed in a specific set of activated or repressed exons by a given splicing factor) were kept. The mutation procedure was stopped when *E* was reached. Second, exonic sequences (MUT exons), selected by sampling human coding exons, were mutated using the same principle used for CUB sequences. In each case, 100 sets of 300 exonic sequences with specific features were generated. A t-test was performed to compare the frequency of amino acid physicochemical properties between the generated sets.

Exonic sequences encoding for specific amino acid physicochemical properties were generated from sampled human coding exons (with the same criteria as CUBs). These sequences were modified by codon substitution to increase the frequency of amino acids encoding for a given physicochemical property P1 and to decrease the frequency of another given physicochemical property P2. Codons that encode P2 were substituted toward codons encoding P1 following the human codon usage bias. SRSF2-like encoded properties exons were generated using the frequency of very small (0.27) and large (0.34) amino acids measured in SRSF2-activated exons or the frequency of very small (0.21) and large (0.38) amino acids measured in SRSF2-repressed exons. TRA2-like encoded properties exons were generated using the frequency of hydrophilic (0.4) and hydrophobic (0.33) amino acids measured in TRA2-activated exons or the frequency of hydrophilic (0.26) and hydrophobic (0.39) amino acids measured in TRA2-repressed exons. SRSF3-like encoded properties exons were generated using the frequency of hydro-neutral (0.38) and charged (0.17) amino acids measured in SRSF3-activated exons or the frequency of hydro-neutral (0.31) and charged (0.22) amino acids measured in SRSF3-repressed exons. The same procedure as for CUBs was used to compare the frequencies of nucleotides or dinucleotides with a t-test.

### Density charts

For each exon, the frequency of each nucleotide or each amino acid physicochemical property was calculated using the formula (1). The exonic sequences were parsed using a sliding window (of size 1 and step 1). Truncated codons (at 3’ or 5’ exon extremities) or codons downstream of stop codons were ignored. Frequency histograms were then computed with R software.

## Acknowledgments

We gratefully acknowledge support from the PSMN (Pôle Scientifique de Modélisation Numérique) of the ENS de Lyon for the computing resources. We thank the members of the LBMC biocomputing center for their involvement in this project. This work was funded by Fondation ARC (PGA120140200853), INCa (2014-154), ANR (CHROTOPAS), AFM-Téléthon, and LNCC. J.B.C was supported by Fondation de France.

